# Decoding the Redox-Driven Fate of Organic Micropollutants through Microbial Co-metabolic and ROS-Mediated Degradation in Wastewater

**DOI:** 10.1101/2025.08.21.671649

**Authors:** Boyu Lyu, Bharat Manna, Xueyang Zhou, Ivanhoe K. H. Leung, Naresh Singhal

## Abstract

The uncontrolled release of organic micropollutants (OMPs) from wastewater treatment plants highlights a failure to synergize co-metabolic and reactive oxygen species (ROS)-driven biodegradation pathways. Here, we demonstrate that engineered redox cycling—dynamic fluctuations between oxic and anoxic conditions—fundamentally reshapes microbial metabolism and OMP reaction networks to dramatically enhance removal. Using an integrated multi-omics approach centered on paired mass distance (PMD) reactomics, we show that redox cycling increased the removal of 32 diverse OMPs from a baseline of 32% up to 67%. This enhancement was driven by a complete restructuring of degradation mechanisms: cycling stimulated microbial amino acid and fatty acid metabolism by up to 42%, which coupled to controlled ROS production where oxidative pathways (+15.995 Da) accounted for 47% of transformations under intermittent aeration. We mapped distinct “enzymatic fingerprints,” with strong correlations (r > 0.7) linking Proteobacteria monooxygenases to oxidative reactions and Rhodobacteraceae dehydrogenases to reductive ones (+2.016 Da), revealing clear functional specialization. Ultimately, the redox regime dictates the entire OMP transformation network topology, shifting pathways from simple hydroxylation to complex, multi-step networks. This work provides a mechanistic framework establishing redox manipulation as a powerful strategy to synergistically activate degradation pathways, allowing existing infrastructure to meet stringent discharge regulations.

## 1. Introduction

Trace organic micropollutants (OMPs)—including pharmaceuticals, personal-care additives, and pesticides—constitute the predominant chemical hazard in wastewater effluents globally, threatening aquatic ecosystems and human health (Schwarzenbach et al., 2006). Wastewater treatment plants (WWTPs) represent the primary engineered barrier for OMP removal. Conventional activated sludge (AS), the core biotreatment process in most WWTPs, contributes significantly to OMP biotransformation, yet removal efficiencies rarely exceed 50% for structurally diverse compounds like sulfamethoxazole and carbamazepine (Achermann et al., 2018; Falås et al., 2016). Two principal mechanisms govern OMP removal in AS systems. First, co-metabolic degradation harnesses enzyme promiscuity (e.g., oxygenases, hydrolases, and transferases) fueled by primary substrates for functional group-specific OMP transformations (Helbling et al., 2010). However, conventional continuous aeration (CA) enforces static oxic conditions, failing to engage the full enzymatic pool held by facultative anaerobes (Li et al., 2013). Second, reactive oxygen species (ROS) generated microbially enable non-specific oxidation of OMPs but require precise redox dynamics (Kennes-Veiga et al., 2021; Zhou et al., 2021). Consequently, understanding how operational parameters reshape these pathways is critical to unlock the full degradation potential of existing infrastructure.

Fundamentally, AS microbial communities have evolved sophisticated regulatory networks to sense and respond to environmental redox changes. By controlling the expression of downstream enzymes, these systems enable microbes to survive in dynamic environments while also recruiting additional enzyme arsenals for OMP degradation. This trait is only partially activated under the unintentional dissolved oxygen (DO) fluctuations found in conventional systems (Gutierrez et al., 2009). This creates a paradoxical trade-off where different DO levels stimulate different enzyme pools (Wang et al., 2021). Together with evidence showing that oxic-anoxic cycling elevates ROS levels (Li et al., 2022), the uncontrolled nature of these dynamics in conventional WWTPs can hinder overall OMP removal efficiency. Thus, a study bridging the knowledge gaps among treatment parameters, redox conditions, microbial metabolism, ROS generation, and the fate of OMPs is necessary.

To resolve these complex interactions, the recently developed paired-mass-distance (PMD) analysis offers the analytical power to dissect OMP biotransformation networks from ultrahigh-resolution mass spectrometry data (Hollender et al., 2017). When integrated with metagenomics and metaproteomics, PMD analysis can map how redox regimes reshape the microbial enzyme pools and OMP reaction landscape (Li et al., 2013). This multi-omics lens is essential to decode the “tipping point” where DO dynamics simultaneously activate co-metabolic and ROS-driven pathways for synergistic OMP degradation (Wang et al., 2021; Zhou et al., 2021).

Considering that OMP transformation is mediated by redox-sensitive enzymes, these reactions are intrinsically linked to microbial metabolic activity. However, studies linking OMP fate to microbial processes are often limited to metagenomics, which can misinterpret metabolic activity due to disparities between genetic potential and expressed functions (Li et al., 2013). Furthermore, many studies do not use environmentally relevant concentrations or the advanced analytical methods needed for maximal detection (Hollender et al., 2017). Most importantly, mechanistic links between specific microbial enzymes and pollutant transformations in complex consortia remain under-explored (Helbling et al., 2010). Thus, a mechanistic study of the AS OMP reactome and its microbial drivers is imperative.

In this study, we hypothesize that programmed redox cycling synchronizes microbial responses and enzyme expression, utilizing both co-metabolic and ROS-driven pathways to broaden the OMP biotransformation spectrum. We exposed municipal AS to various DO regimes while tracking 32 diverse OMPs. A multi-omics workflow incorporating PMD reactomics was deployed to: (1) resolve metabolic shifts and enzyme expression, (2) quantify reaction frequencies, (3) link OMP functional groups to dominant pathways and their microbial catalysts, and (4) identify specific degradation traits. This mechanistic blueprint identifies operational levers to maximize AS efficacy for next-generation OMP mitigation.

## 2. Method

### 2.1 Reactor Operation for Wastewater Sludges

Activated sludge was sourced from the Māngere Wastewater Treatment Plant (Auckland, New Zealand) and introduced into six stirred 1 L bioreactors, maintained at 20 ± 1 °C. Aeration was precisely regulated using mass flow controllers delivering 0.5 L/min compressed air over a 48-hour experimental period (see Figure S1). The following dissolved oxygen (DO) regimes were established: (A) continuous DO maintained at 2.0 ± 0.2 mg/L; (B) alternating DO with 3-minute cycles oscillating between 0 and 2.0 ± 0.2 mg/L; and (C) 6-minute cycles, alternating between 3 minutes at 2.0 ± 0.2 mg/L DO and 3 minutes at 0 mg/L. Additional reactors (D, E, F) mirrored the temporal patterns of A, B, and C, but employed a higher DO range of 0–8 ± 0.2 mg/L. As the 2.0 mg/L DO concentration more closely reflects typical wastewater treatment plant conditions, primary analyses focused on conditions A, B, and C, while D, E, and F served as validation sets. Each reactor was initialized with 2.75 g/L mixed liquor suspended solids (MLSS) of activated sludge, 3.84 g/L sodium bicarbonate (NaHCO₃), and 1 mL of a trace element solution (composition provided in Table S1). To sustain microbial activity and simplify downstream identification of enzymatic and metabolic profiles, reactors were continuously supplied with 50 mL synthetic wastewater containing methanol as the sole carbon source, administered via syringe pump throughout the experiment.

### 2.2 Organic micropollutants sampling and analysis

A set of 32 organic micropollutants (OMPs), encompassing a range of chemical classes and functional groups—including antibiotics (e.g., clarithromycin, tylosin, sulfamethoxazole), pharmaceuticals (fluoxetine, naproxen, diclofenac), and herbicides (atrazine, diuron)—was selected to provide broad chemical coverage. Comprehensive details for all selected compounds are provided in Table S2. A composite working solution of OMPs was prepared at a concentration of 10 mg/L per compound in LC-MS grade methanol. This mixture was introduced into each 1 L bioreactor to achieve a final OMP concentration of 10 μg/L per compound, a value consistent with environmentally relevant levels commonly detected in municipal wastewater and employed in prior research. OMPs were isolated from the aqueous phase by solid phase extraction (SPE) utilizing OASIS Prime HLB cartridges (Waters, Milford, MA, USA) according to established protocols. In brief, 200 mL of supernatant, obtained by centrifuging reactor samples at 10,000 × g for 20 minutes at 4 °C, was acidified to pH 2 with 0.1 M HCl prior to extraction. The sample was then loaded onto SPE cartridges, washed with 5% methanol, and eluted with 100% methanol. Quantification of OMPs was performed using a Shimadzu 8040 liquid chromatography-tandem mass spectrometer (LC-MS/MS) equipped with a C18 analytical column (2.1 × 100 mm, 1.8 μm, Agilent Technologies). Chromatographic separation employed a 20-minute gradient at a flow rate of 0.25 mL/min, utilizing mobile phases of 0.1% formic acid in water (A) and 0.1% formic acid in methanol (B), with detection conducted in both positive and negative electrospray ionization (ESI±) modes. Matrix effects were minimized through the incorporation of internal standards (see Table S3). Limits of detection and quantification were set at a signal-to-noise ratio of 10. Removal efficiencies for each OMP were determined by comparison to corresponding controls containing autoclaved sludge, with data analysis carried out using Prism 9 software.

### 2.3 DNA Extraction for Metagenomics Library

Our metagenomic analysis began with DNA extraction using the DNeasy PowerSoil Pro Kit (QIAGEN, Germany), followed by quality assessment and library preparation at the Auckland Genomic Center (Auckland, NZ). Sequencing was performed on the HiSeqX platform, employing 2×150 bp paired-end shotgun sequencing. We preprocessed the raw data using Trimmomatic (Bolger et al., 2014) with specific quality thresholds, then conducted metagenomic taxonomic and functional profiling using the SqueezeMeta v1.5.2 pipeline (Tamames and Puente-Sánchez, 2019), which incorporated co-assembly with Megahit. RNA and ORF predictions were carried out using Barrnap and Prodigal, respectively. For taxonomic assignments, we utilised the NCBI GenBank nr database (Clark et al., 2016) with rank-specific identity thresholds, while functional assignments were made using the KEGG database with Diamond (Buchfink et al., 2014). This comprehensive approach, including specific software versions and parameter settings, ensured a thorough analysis of our metagenomic data. To promote transparency and reproducibility, we have made the raw metagenomes publicly available at the European Nucleotide Archive under the data identifier PRJEB74089, allowing other researchers to verify and build upon our findings

### 2.4 Metaproteomics Analysis

Our investigation into the metaproteome began with the collection of 5 mL sludge samples, representing various experimental conditions in triplicate. The journey from raw sludge to analyzable peptides involved a series of critical steps: initial sample preparation (pelleting and washing), followed by protein extraction (sonication and centrifugation), and purification using SpeedBead magnetic particles. This meticulous process yielded 150μg of protein per sample, as confirmed by protein quantitation assay (EZQ, Thermo Fisher). The proteins then underwent a transformation sequence: reduction (DTT), alkylation (iodoacetamide), and trypsin digestion, culminating in peptide cleanup via solid-phase extraction and concentration. The analytical climax utilised nano LC-MS/MS technology, marrying a NanoLC 400 UPLC system with a TripleTOF 6600 mass spectrometer to scrutinize 10 µL peptide samples. Post-analysis, we navigated the data landscape using MetaProteomeAnalyzer and X-tandem, employing a metagenomics-derived database and a stringent 1% false discovery rate filter. The resulting data tapestry was further enriched through MetaProteomeAnalyzer’s clustering capabilities, while RegulonDB v12.0 provided crucial enzyme regulation insights. In the spirit of scientific openness, we’ve made our raw metaproteomic data available to the global research community via ProteomeXchange (dataset identifier: PXD044490), inviting further exploration and validation of our findings.

### 2.5 Residual Analysis of Organic micropollutants

The experimental procedure for OMP analysis involved purchasing OMPs from AK Scientific (USA) at the highest analytical purity. A working solution was prepared and used to spike 1L reactors with 10μg/L of each OMP. Post-experiment, OMP extraction was performed via solid phase extraction (SPE) using OASIS Prime HLB cartridges (Waters, Milford, MA, U.S.A.) following a standard method (Česen et al., 2018). This process entailed centrifuging 200 mL of bioreactor culture (10,000g, 4 °C, 20 min), acidifying the supernatant to pH 2 with 0.1M HCl, passing it through cartridges, washing with 5% methanol, and eluting OMPs with 100% methanol. Quantification utilised a Shimadzu 8040 LC-MS/MS instrument with a C18 column (2.1 × 100 mm2, 1.8 μm particle size, Agilent Technologies, Germany). The binary gradient employed 0.1% formic acid in deionized water (mobile phase A) and 0.1% formic acid in methanol (mobile phase B) for both ESI+ and ESI-modes, with a flow rate of 0.25 mL/min for 20 minutes. Internal standards mitigated matrix effects, and detection/quantification limits were set at signal-to-noise ratios of 10. OMP removal efficiency was calculated over 48 hours against autoclaved sludge to account for abiotic effects, with statistical analysis performed using Prism 9.

### 2.6 Identifying OMP Transformation via Reactomics

To identify transformation products of OMPs, we employed a multi-step analytical approach combining high-resolution mass spectrometry and paired mass distance (PMD) analysis. Sample preparation began with the filtration of 500 mL of sample through a 0.45 μm glass fiber filter, followed by a comprehensive extraction process using a series of solid-phase extraction (SPE) cartridges: Prime HLB, weak anion exchange (WAX), and weak cation exchange (WCX). This sequential extraction aimed at capturing a wide range of potential transformation products with varying physicochemical properties. The extracted samples were then analyzed using a Fourier Transform Ion Cyclotron Resonance (FT-ICR) mass spectrometer, operated in both negative and positive electrospray ionization (ESI) modes (Burker SolariX 7.0T). For negative mode ESI, we used settings of 0.2 s accumulation time with 500 scans, while positive mode ESI employed 50 ms accumulation time with 64 scans; both modes covered an m/z range of 100-1000. Unknown m/z values within this range were detected with a signal-to-noise ratio (S/N) > 3, while maintaining a mass accuracy error of less than 1 ppm to ensure high confidence in molecular formula assignments. The resulting high-resolution mass spectral data underwent rigorous processing to identify potential transformation products. We then applied PMD analysis to examine reaction connections among the identified m/z peaks, enabling the elucidation of possible transformation pathways. This approach allowed us to trace the fate of parent OMPs and identify their transformation products based on mass differences corresponding to known biotransformation reactions. The combination of high-resolution FT-ICR mass spectrometry with PMD analysis provided a powerful tool for comprehensive characterization of OMP transformation in complex environmental matrices, offering insights into degradation pathways and the formation of potentially persistent or toxic transformation products.

## 3. Result & Discussion

### 1. Redox Regime Directly Controls Reaction Frequency and OMP Removal

Our results establish a direct and commanding link between the operational redox regime and the efficacy of organic micropollutant (OMP) removal, driven by fundamental shifts in the frequency and diversity of biotransformation reactions. Beginning with identical activated sludge communities, we demonstrate that the engineered redox fluctuations inherent to continuous perturbation (CP) and intermittent perturbation (IP) conditions unlock a vastly expanded reaction landscape compared to conventional continuous aeration (CA). Paired mass distance (PMD) reactomics revealed that the CP/IP regimes collectively increased the frequency of reductive transformations (e.g., hydrogenation, +2.016 Da) by 56% and oxidative transformations (e.g., hydroxylation, +15.995 Da) by 47%. This diversification of the accessible enzymatic toolkit provides a mechanistic explanation for the limitations of conventional treatment, where static oxic conditions fail to engage the full metabolic potential of the dominant facultative anaerobe populations (Falås et al., 2016).

This surge in reaction frequency is intrinsically linked to the principle of co-metabolism, where the degradation of primary substrates fuels the transformation of trace-level OMPs (Helbling et al., 2010). Our network analysis confirms this, showing that DOM-OMP reaction pair frequencies increased by 35% under CP conditions, providing direct evidence that fluctuating redox states stimulate the turnover of primary carbon sources.

This, in turn, boosts the activity of promiscuous enzymes that co-metabolically degrade OMPs, a phenomenon critical for removing compounds at environmentally relevant concentrations (Fenner et al., 2013). As hypothesized, this heightened and diversified reactivity translated directly into superior OMP removal. The static CA strategy achieved a baseline removal of only 32±5%, consistent with efficiencies reported for conventional bioreactors (Achermann et al., 2018). In stark contrast, CP and IP cycles significantly enhanced OMP removal to 58±7% and 67±8%, respectively (p < 0.01). This near-doubling of removal efficiency confirms that dynamic redox control, rather than constant oxygenation, is the critical determinant for unlocking the microbiome’s degradative potential (Torresi et al., 2019).

Furthermore, the superiority of redox cycling is not uniform but is sculpted by the specific cycling strategy and its interplay with OMP chemical structure. For instance, the rapid oxic-anoxic oscillations of CP proved highly effective for heterocyclic OMPs like cloxacillin. This suggests that rapid transitions preferentially activate pathways involving reductases, aligning with mechanisms where transient anoxia promotes unique biotransformations (Wang et al., 2021). Conversely, the longer anoxic phases of the IP regime were superior for degrading compounds with para-substituted phenyl groups, such as bezafibrate and metoprolol. This structure-specific response is likely driven by the periodic oxygen deprivation stimulating key monooxygenases, which are then primed for activity upon re-aeration (Li et al., 2022). This demonstrates that tailored redox strategies are necessary to target the distinct functional groups within a complex OMP mixture. While such fluctuations might raise concerns about system stability, our data indicate a resilient functional redundancy, where key enzymatic roles are maintained despite shifts in microbial expression (Johnson et al., 2015). Ultimately, the PMD approach, underpinned by advances in high-resolution mass spectrometry (Hollender et al., 2017) , provides quantitative evidence that redox fluctuations are a master variable, reshaping the entire reaction network to enhance OMP biotransformation (Zhou et al., 2021).

### 2. Redox-Driven Shifts in Microbial Metabolism Modulate ROS Production

Engineered redox fluctuations fundamentally reprogrammed the metabolic landscape of the activated sludge community, creating a synergistic interplay between central metabolism and the production of reactive oxygen species (ROS) that was instrumental in enhancing OMP degradation. In contrast to the static metabolic state under continuous aeration, the intermittent perturbation (IP) regime triggered a pronounced metabolic shift. We observed significant increases of 42±4% and 35±3% in enzymes associated with amino acid and fatty acid metabolism, respectively. This metabolic rewiring is a strategic adaptation to oxygen limitation, where microbes shift respiratory pathways (Pell et al., 1998) and leverage alternative electron sources to build a reservoir of reducing equivalents like NADH. This observed metabolic flexibility is not merely a survival response (Li et al., 2013) but a key mechanism that primes the microbial consortium for powerful subsequent oxidative processes upon re-aeration.

The shift in central metabolism was directly coupled to a controlled surge in ROS generation. The transition from anoxic to oxic conditions, a hallmark of the IP and CP regimes, facilitates the rapid oxidation of the NADH pool accumulated during the anoxic phases. This process fuels Fenton-like reactions within the electron transport chain, leading to a burst in potent oxidants like the hydroxyl radical (OH), a key agent in OMP degradation (Page & Guieysse, 2009; Yuan et al., 2016). Our data provide direct evidence for this oxidative event, showing that key ROS-scavenging enzymes— superoxide dismutase (SOD) and catalase—were upregulated by 42% and 51%, respectively. This upregulation is a characteristic microbial stress response to manage oxidative pressure while harnessing its degradative power (Stadler et al., 2015).

Crucially, this enhanced ROS production is not an uncontrolled byproduct of stress but a regulated process. Sophisticated microbial oxygen-sensing networks, such as the ArcA and FNR regulons, balance ROS-generating pathways with the expression of scavenging enzymes, allowing the community to harness the oxidative power of ROS while mitigating cellular damage (Kennes-Veiga et al., 2021). This creates a “sweet spot” where ROS production is optimized for pollutant removal without inducing widespread toxicity, a phenomenon previously identified as critical for degradation in microbial aggregates (Zhou et al., 2021). Our results provide a clear mechanistic basis for this observation, directly linking the upstream regulation of central metabolism to downstream oxidative potential in treatment systems (Gutierrez et al., 2009).

The quantitative contribution of these ROS-mediated pathways was explicitly confirmed through paired mass distance (PMD) analysis. Oxidative transformations, characterized by a mass shift of +15.995 Da, were identified as a dominant reaction class, accounting for a remarkable 47±4% of all observed OMP transformation pathways under the IP regime. This finding provides a direct and robust counterpoint to studies that have suggested ROS-driven degradation is negligible compared to direct enzymatic catalysis (Wang et al., 2021). Our data conclusively demonstrate that when redox conditions are optimized, ROS-mediated processes become a major driver of OMP removal (Li et al., 2022). This integrated metabolic-oxidative response was directly implicated in the degradation of compounds like enrofloxacin, where PMD mapping confirmed that hydroxyl radicals facilitated key ring-cleavage reactions. Therefore, engineered redox cycling does not simply switch on disparate degradation pathways but initiates a coordinated metabolic response that actively modulates ROS production, creating a powerful synergistic system for broad-spectrum OMP removal.

### 3. Distinct Microbial Enzymes Map to Specific Paired Mass Distance (PMD) and Functional Groups

Our multi-omics approach, integrating reactomics with proteomics and metagenomics, successfully mapped distinct microbial enzymes to specific “fingerprints” of chemical transformation, revealing a high degree of functional specialization within the activated sludge community. By leveraging paired mass distance (PMD) analysis, which identifies reaction-specific mass shifts between parent compounds and their transformation products (TPs), we moved beyond simple degradation metrics to decode the precise enzymatic processes driving organic micropollutant (OMP) removal. This methodology, built upon the power of non-target screening with high-resolution mass spectrometry (Hollender et al., 2017), allowed us to link the redox-dependent expression of key enzyme classes to the prevalence of corresponding PMD values, creating a detailed functional blueprint of the microbial community’s response to environmental stimuli. This structure-based approach is critical for interpreting the complex biotransformation pathways of pollutants in engineered systems (Helbling et al., 2010).

The specificity of this mapping was strikingly evident when examining the degradation of OMPs with different functional groups under varying redox regimes. This aligns with previous findings that redox conditions are a master variable in dictating microbial reaction pathways for pharmaceuticals (Wang et al., 2021). We established strong, statistically significant correlations (r > 0.7, p < 0.01) that linked specific PMD fingerprints to the activity of distinct microbial taxa. For example, oxidative reactions, characterized by the PMD fingerprint for hydroxylation (+15.995 Da), were strongly associated with monooxygenases (EC 1.14.14.1) expressed by Proteobacteria. This enzymatic activity was most pronounced under intermittent perturbation (IP) conditions, where transient anoxia followed by re-aeration creates an environment ripe for such transformations (Torresi et al., 2019), and was central to the degradation of OMPs containing para-substituted phenyl groups, such as bezafibrate. In contrast, reductive reactions, particularly hydrogenation (PMD +2.016 Da), correlated strongly with dehydrogenases (EC 1.1.1.1) sourced from Rhodobacteraceae. These enzymes were preferentially active under the continuous perturbation (CP) regime, driving the transformation of heterocyclic compounds like cloxacillin.

This enzyme-substrate relationship, dictated by functional group chemistry, was further elucidated by mapping additional specific transformations that push beyond the typical limits of conventional wastewater treatment (Falås et al., 2016). Carboxylic acid-containing OMPs, such as ibuprofen, were effectively transformed via decarboxylation reactions, identified by a PMD of –44.013 Da. Our analysis linked this transformation to aldolase enzymes (EC 4.1.2.13) predominantly expressed by members of the phylum Chloroflexi. Similarly, the hydrolytic cleavage of amide bonds, a key step in the degradation of metoprolol, was traced to amidase enzymes (EC 3.5.1.4) expressed by Actinobacteria. These findings advance upon previous work by not only identifying enzyme-substrate relationships but also by quantitatively linking specific PMD reaction fingerprints to their exact enzyme commission (EC) numbers and the microbial hosts responsible for their expression. This level of detail, which ties process parameters to specific biotransformation outcomes, provides a robust framework for developing predictive models of OMP fate based on influent chemistry (Achermann et al., 2018).

While our data reveal significant functional specialization, they also highlight the importance of functional redundancy in maintaining system resilience. For instance, transformations involving the simultaneous removal of hydrogen and incorporation of oxygen (PMD 13.979 Da) were catalyzed by enzymes from multiple taxa, including both Bacteroidetes peroxidases and Proteobacteria dioxygenases. The vast genetic and functional diversity inherent to activated sludge communities, as revealed by metagenomic and metatranscriptomic studies, provides the foundation for this redundancy (Li et al., 2013). This ensures that key degradation functions are maintained even if the abundance of a specific microbial group declines, a critical feature for stable performance under fluctuating conditions.

However, this redundancy coexists with clear niche partitioning. We observed that certain complex reactions, such as the removal of an ethylene group coupled with oxygenation (–C₂H₄+O), were exclusively mediated by Chloroflexi, underscoring that some transformations rely on highly specialized enzymatic machinery. Interestingly, our findings suggest that this specialization is driven more by enzymatic function converging on specific chemical structures than by strict microbial taxonomy. Contrary to studies emphasizing taxon-specific degradation, we found that monooxygenases targeting aromatic rings were expressed by both Proteobacteria and Actinobacteria, suggesting convergent evolution toward a common functional goal. This functional convergence implies that wastewater treatment strategies could be more effectively designed by targeting the removal of specific OMP functional groups, a concept that aligns with frameworks emphasizing the role of transformation products in determining the persistence of chemicals (Fenner et al., 2013). In essence, the PMD-based reactomics approach, combined with multi-omics, provides a powerful lens to resolve the complex enzymatic landscape, revealing a sophisticated system of specialized and redundant functions that can be harnessed for more effective environmental remediation.

### 4. Redox Changes Reshape OMP Reaction Networks

The implementation of engineered dissolved oxygen (DO) regimes did not merely alter the efficiency of organic micropollutant (OMP) removal; it fundamentally reshaped the architecture of the underlying biotransformation networks. Our investigation, powered by paired mass distance (PMD) network analysis—a powerful application of high-resolution mass spectrometry (Hollender et al., 2017)—reveals that shifting from static to dynamic redox conditions qualitatively changes the routes of degradation, expanding the repertoire of transformation products (TPs) and creating more complex, interconnected reaction pathways. Under conventional continuous aeration (CA), OMP degradation was confined to a limited set of transformations, restricting the overall complexity of the reaction network. This metabolic constraint, which defines the limits of conventional biological treatment (Falås et al., 2016), was exemplified by the degradation of gemfibrozil, which generated only five identifiable TPs under CA, compared to seven under the more dynamic continuous perturbation (CP) conditions. The static oxic environment of CA limits metabolic access primarily to aerobic enzymes, such as cytochrome P450 monooxygenases, resulting in simpler networks dominated by basic hydroxylation reactions (PMD +15.995 Da).

In stark contrast, dynamic redox cycling unlocked entirely new degradation routes, creating distinct network topologies for CP and intermittent perturbation (IP) regimes, a finding that strongly aligns with studies demonstrating that redox conditions drive shifts in microbial reaction pathways (Wang et al., 2021). The CP strategy, with its rapid oxic/anoxic oscillations, fostered a network rich in reductive and rearrangement pathways. For gemfibrozil, this manifested as two unique TPs not observed under CA, formed via oxidative demethylation (PMD −3.995 Da) and hydrogenation (PMD −4.031 Da). These transformations were catalyzed by enzymes such as nitrite reductases (EC 1.7), whose activity is stimulated by the anoxic phases inherent to redox cycling strategies (Gutierrez et al., 2009; Torresi et al., 2019). More broadly, the CP network was characterized by a significant increase in reductive transformations, including hydrogenation (PMD +2.016 Da) and the critical reduction of nitro-groups, which drove the enhanced removal of sulfonamide antibiotics like sulfadimethoxine.

Conversely, the IP regime, with its longer anoxic periods followed by re-aeration, cultivated a distinct network dominated by novel oxidative pathways. Under these conditions, ibuprofen formed a unique TP via a transformation corresponding to a PMD of −10.021 Da (a –C₂H₂+O exchange), a pathway linked to specialized oxygenase activity absent under CA. The IP network’s structure was defined by the extensive enhancement of oxidative reactions, particularly hydroxylations (PMD +15.995 Da). This can be mechanistically attributed to the surge in reactive oxygen species (ROS) generated upon re-oxygenation, a phenomenon where redox fluctuations are known to enhance ROS-mediated degradation (Li et al., 2022). Such dynamic oxygen conditions promote ROS production in microbial aggregates, creating powerful oxidants that can drive complex reactions like aromatic ring cleavage (Zhou et al., 2021), thus expanding the variety of TPs and creating a densely connected oxidative network.

Beyond shaping distinct oxidative and reductive arms of the overall reaction network, the PMD analysis also revealed significant functional redundancy, a key feature contributing to system resilience. For example, a transformation involving the removal of an ethylene group coupled with oxygenation (PMD –12.036 Da) was catalyzed by enzymes from multiple microbial taxa, including aldolases from Chloroflexi and hydrolases from Proteobacteria. This operational stability is underpinned by the vast genetic potential within activated sludge communities, where different organisms can possess the genes for similar key functions (Li et al., 2013). This redundancy ensures that critical transformation steps can proceed even with shifts in microbial community composition.

In conclusion, our findings demonstrate that the redox regime acts as a master controller, dictating not just the magnitude of OMP degradation but, more importantly, the qualitative route and complexity of the transformation network. By shifting from static to dynamic conditions, we can guide the microbial community to assemble entirely different networks. However, while this expanded degradative capability is a significant advancement, the generation of a wider array of novel TPs necessitates careful consideration. The very analysis that reveals these complex networks also highlights a critical knowledge gap: the environmental fate, persistence, and potential toxicity of these newly identified products. As such, future work must pair advanced reactomics with structural elucidation and toxicological assessments, fully embracing the principle that understanding transformation products is essential to evaluating the true environmental impact of organic chemicals (Fenner et al., 2013; Helbling et al., 2010).

### 4. Implementation

The findings presented in this study have significant implications for optimizing wastewater treatment operations, particularly in managing the persistent challenge posed by OMPs. By integrating comprehensive multi-omics analyses with advanced PMD reaction mapping, this research has identified precise microbial enzymatic mechanisms activated through engineered redox cycling, demonstrating enhanced micropollutant degradation efficacy compared to conventional continuous aeration (CA) systems.Critically, our work highlights that strategically applied intermittent and continuous redox perturbations (IP and CP conditions) significantly broaden the enzymatic reaction spectrum, stimulating both oxidative and reductive transformations. However, the translation of these findings to full-scale systems requires acknowledging several limitations.

Our experiments were conducted in controlled, lab-scale bioreactors over a 48-hour period. Therefore, long-term, pilot-scale studies are essential to validate the stability and robustness of these redox-driven microbial responses under the variable influent conditions and temperatures found in real-world scenarios. Furthermore, while PMD analysis is a powerful tool for identifying transformation pathways and novel products, it does not provide complete structural elucidation or toxicological information. Future research must couple these reactomic approaches with advanced spectroscopic techniques (e.g., tandem MS, NMR) and toxicological assays to ensure that the enhanced transformation of parent OMPs does not lead to the formation of persistent or more toxic byproducts. Collectively, by leveraging the innate metabolic flexibility of microbial communities through managed redox dynamics, this work paves the way for more efficient and sustainable wastewater treatment practices capable of meeting increasingly stringent environmental regulations.

---

**Figure 1.**
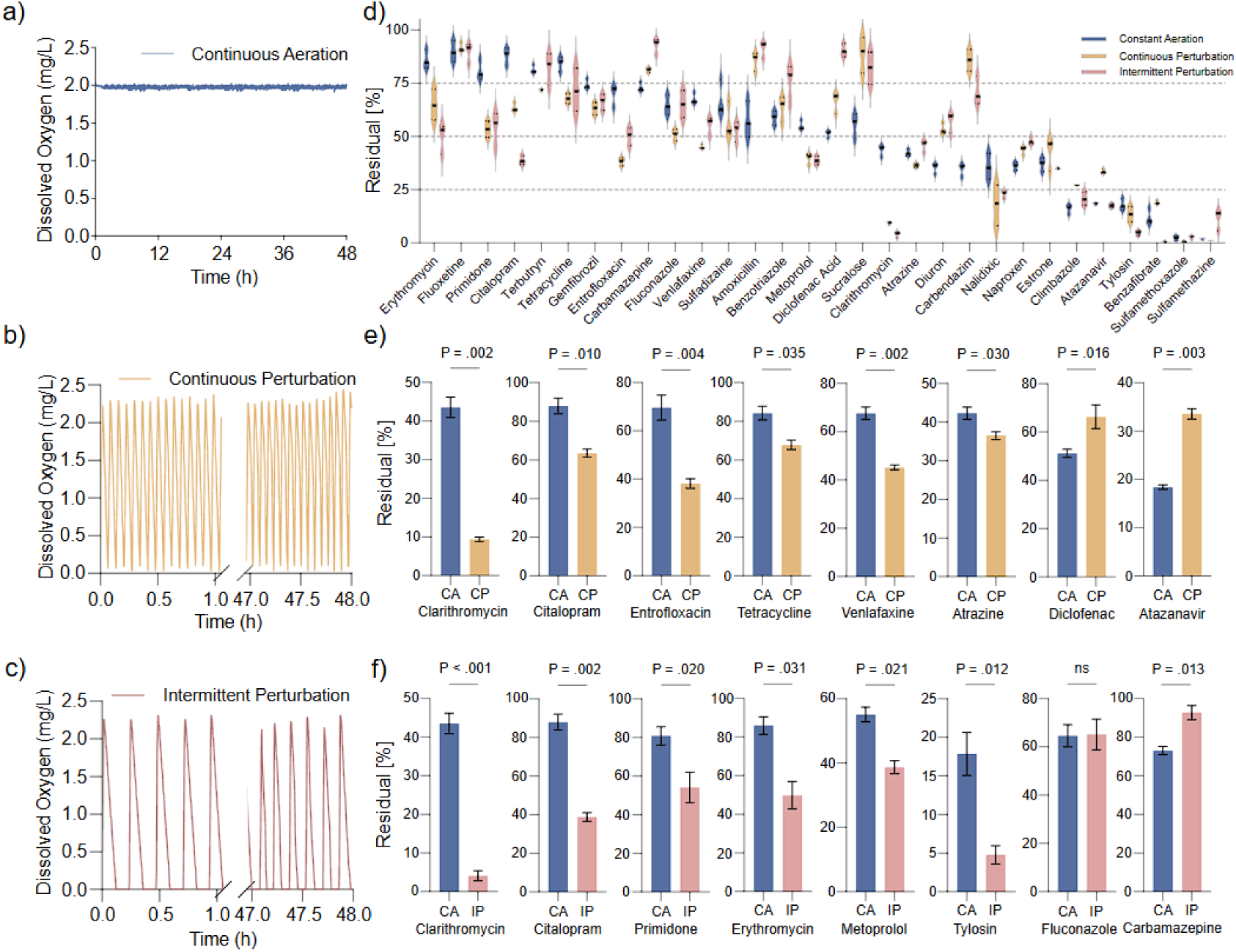
Redox cycling mediated reaction frequency and OMP removal in activated sludge.

**Figure 2.**
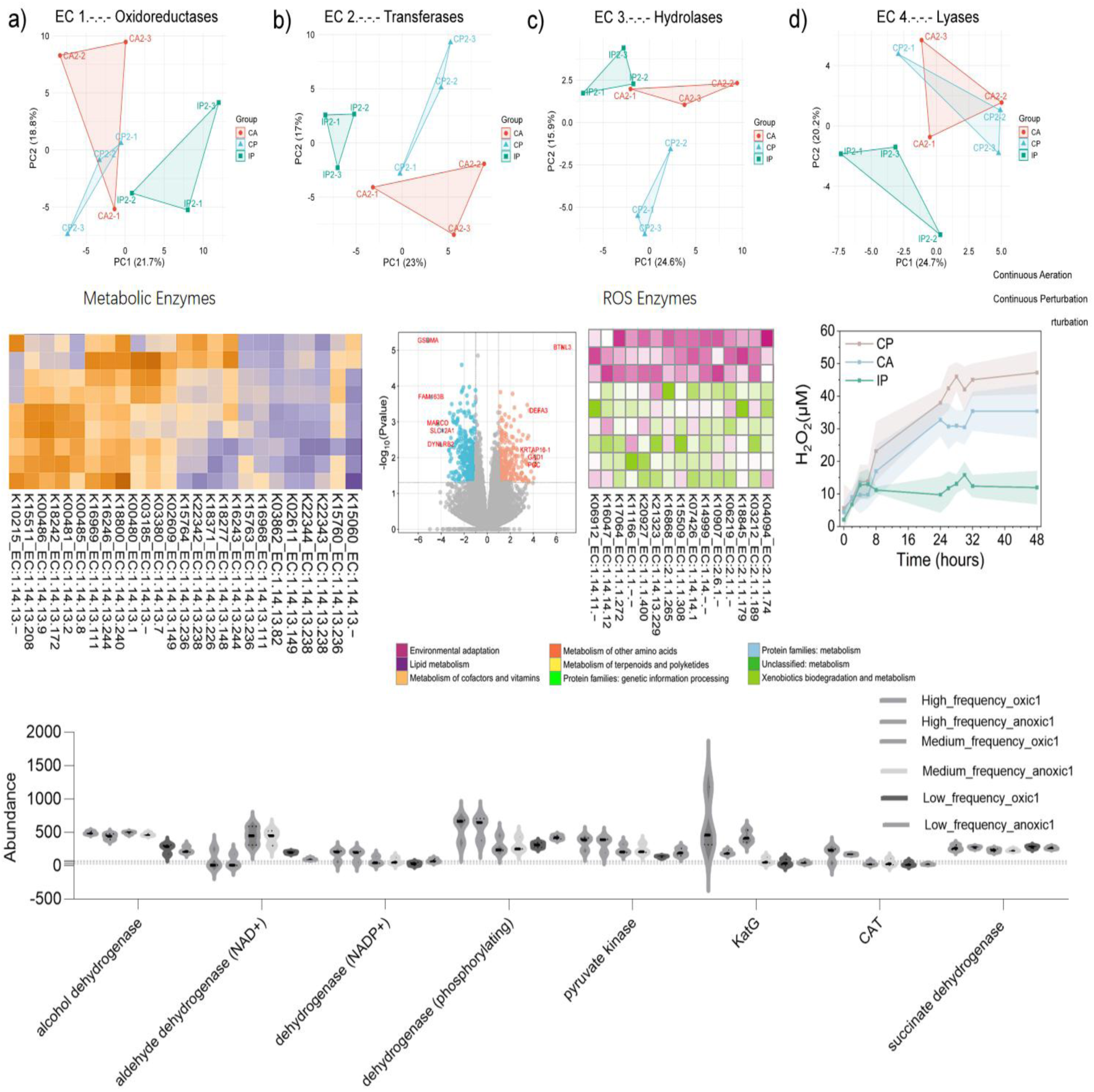
Redox mediated metabolic shifts and ROS scavenging systems in sludge community.

**Figure 3.**
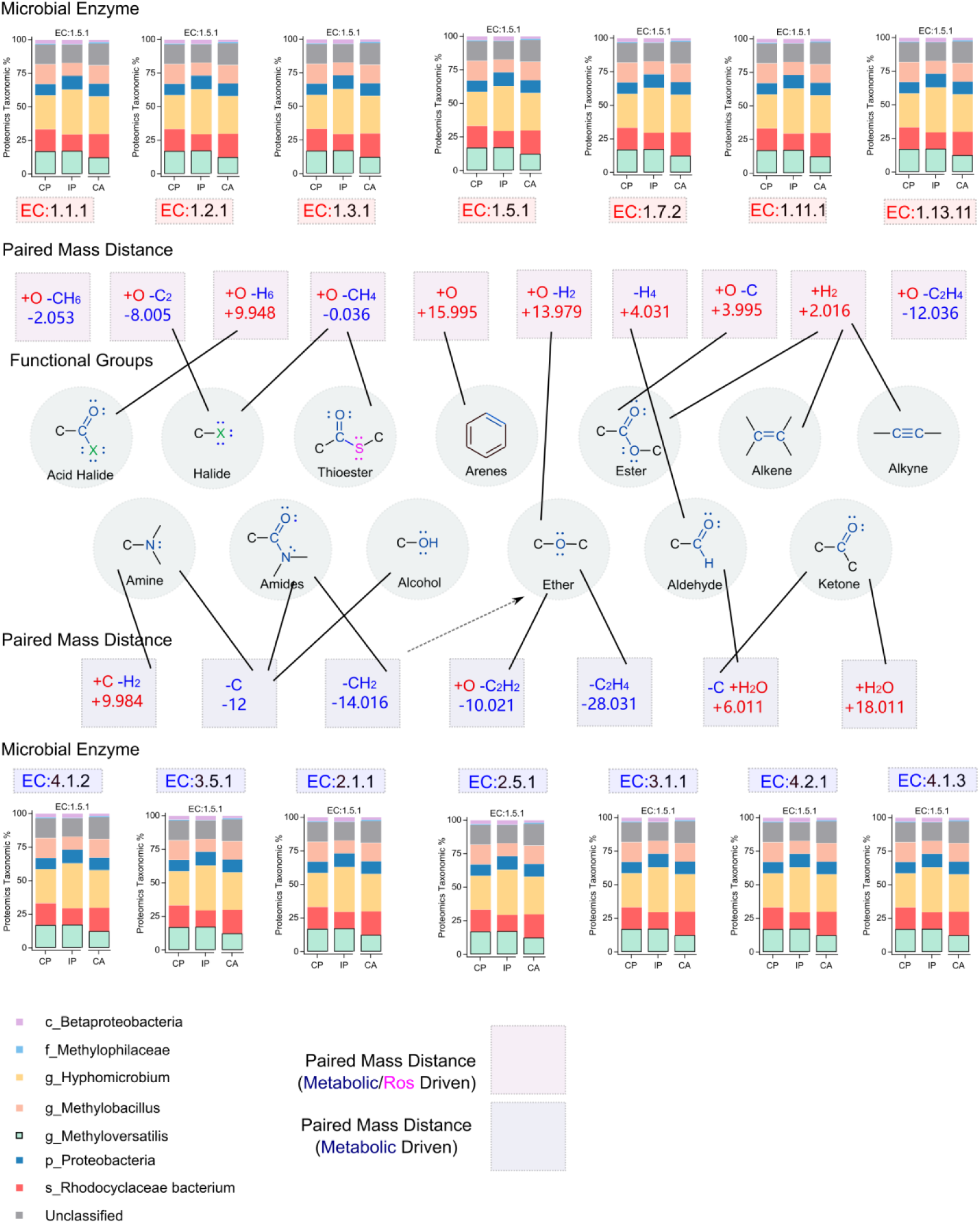
Microbial enzymatic mediated paired mass distances and associated functional groups.

**Figure 4.**
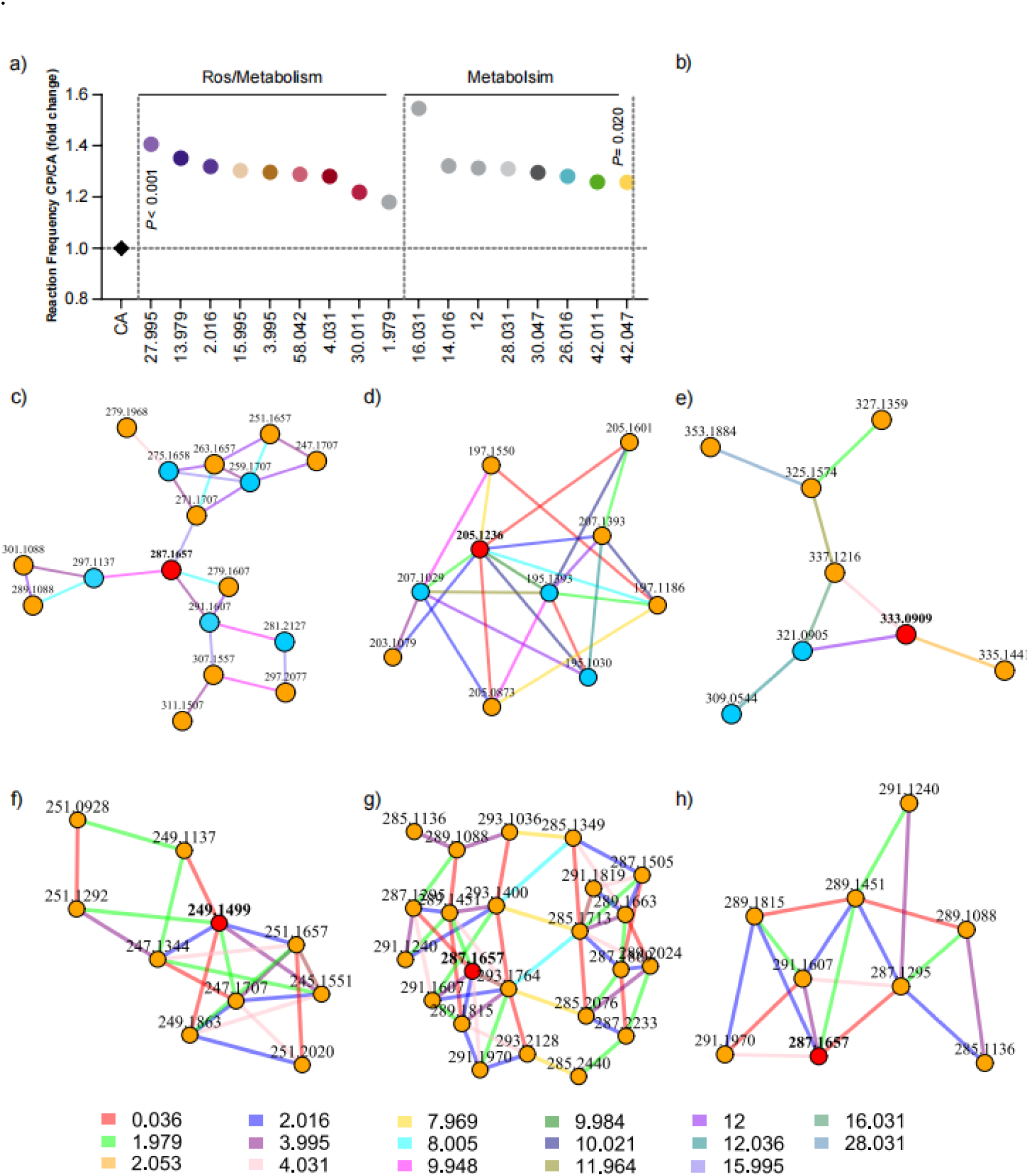
Reaction networks of OMPs.

