## Supplemental Material for "Decoding the Redox-Driven Fate of Organic Micropollutants through Microbial Co-metabolic and ROS-Mediated Degradation in Wastewater"

Total Pages:

Text:

Tables:

Figures:

**Supplementary Tables**

**Table S1** Trace elements and synthetic wastewater provided to the reactor

| **Chemical** | **Concentration (g/L)** |
| --- | --- |
| EDTA | 2.50 |
| ZnSO_4_·7H_2_O | 1.10 |
| CoCl_2_·6H_2_O | 0.80 |
| MnCl_2_·4H_2_O | 2.55 |
| MgSO_4_·7H_2_O | 20.0 |
| CuSO_4_·5H_2_O | 0.86 |
| (NH_4_)_6_Mo_7_O_24_·4H_2_O | 0.07 |
| CaCl_2_·2H_2_O | 2.75 |
| FeSO_4_·7H_2_O | 2.57 |
| **Synthetic wastewater** | **Concentration** |
| Methanol (mL/L) | 26.94 |
| NH_4_Cl (g/L) | 24.46 (C); 30.57 (B & A) |
| KH_2_PO_4_ (g/L) | 2.76 |
| K_2_HPO_4_ (g/L) | 2.76 |
| NaHCO_3_ (g/L) | 76.80 |

**Table S3.** Internal Standards used in this study

| **Chemical** | Formula |
| --- | --- |
| Acesulfame Potassium-^13^C_4_ | ^13^C_4_H_4_KNO_4_S |
| Atenolol-d_7_ | C₁₄H₁₅D₇N₂O₃ |
| Bezafibrate-d_4_ | C₁₉H₁₆D₄ClNO₄ |
| Clarithromycin-N-methyl-^13^C, D_3_ | C₃₇¹³CH₆₆D₃NO₁₃ |
| Diclofenac-D_4_ | C_14_H_7_D_4_Cl_2_NO_2_ |
| Fluconazole-D_4_ | C_13_H_8_D_4_F_2_N_6_O |
| Fluoxetine-D_5_ | C_17_D_5_H_13_F_3_NO |
| Metoprolol-D_7_ | C_15_H_18_D_7_NO_3_ |
| Sucralose-D_6_  Sulfamethoxazole-^13^C_6_ | C_12_H_13_D_6_Cl_3_O_8_  C_10_H_11_N_3_O_3_S |

**Supplementary Figures**

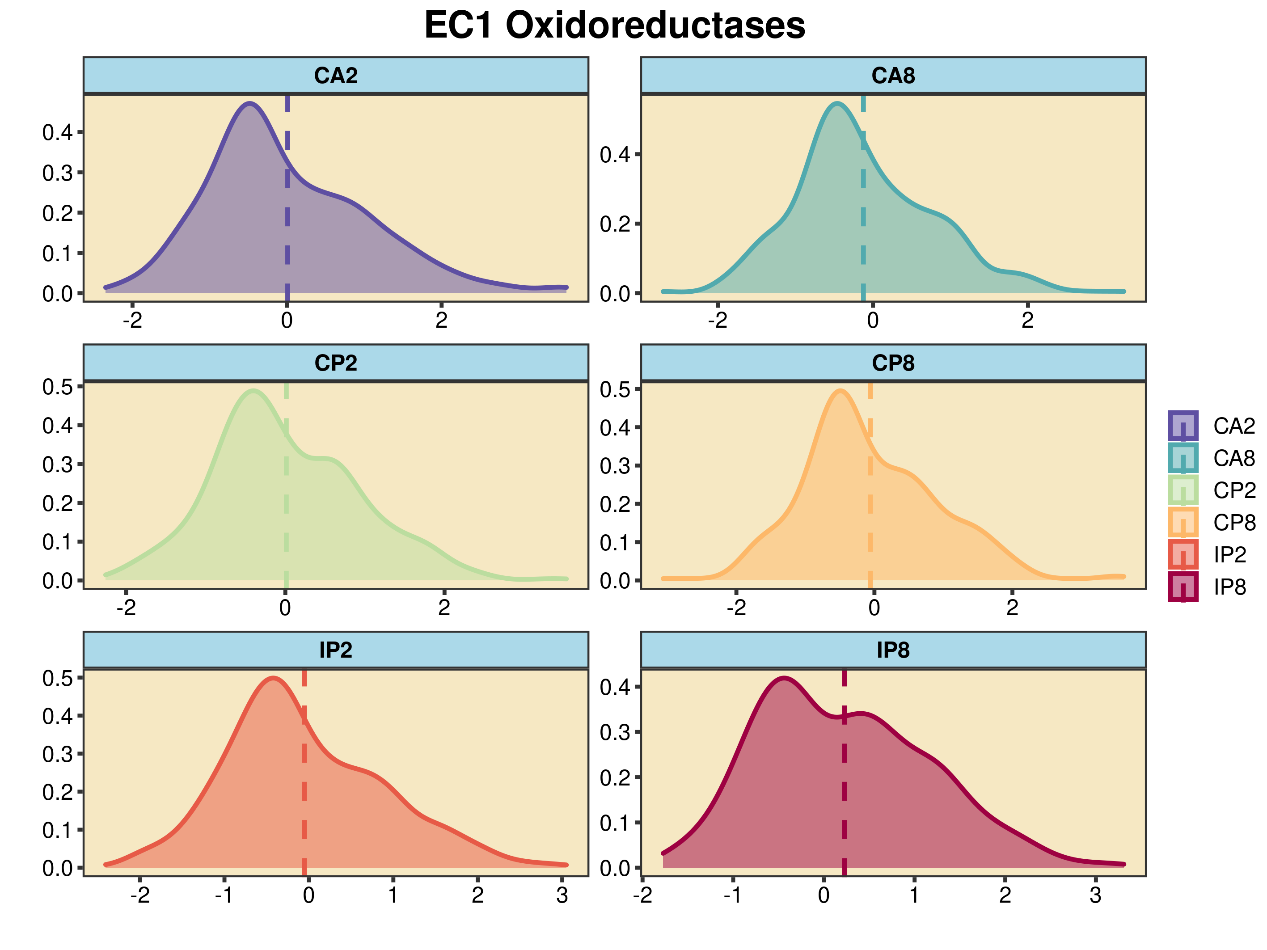

**Figure S1. Distribution and abundance of oxidore4tactase among test conditions.**

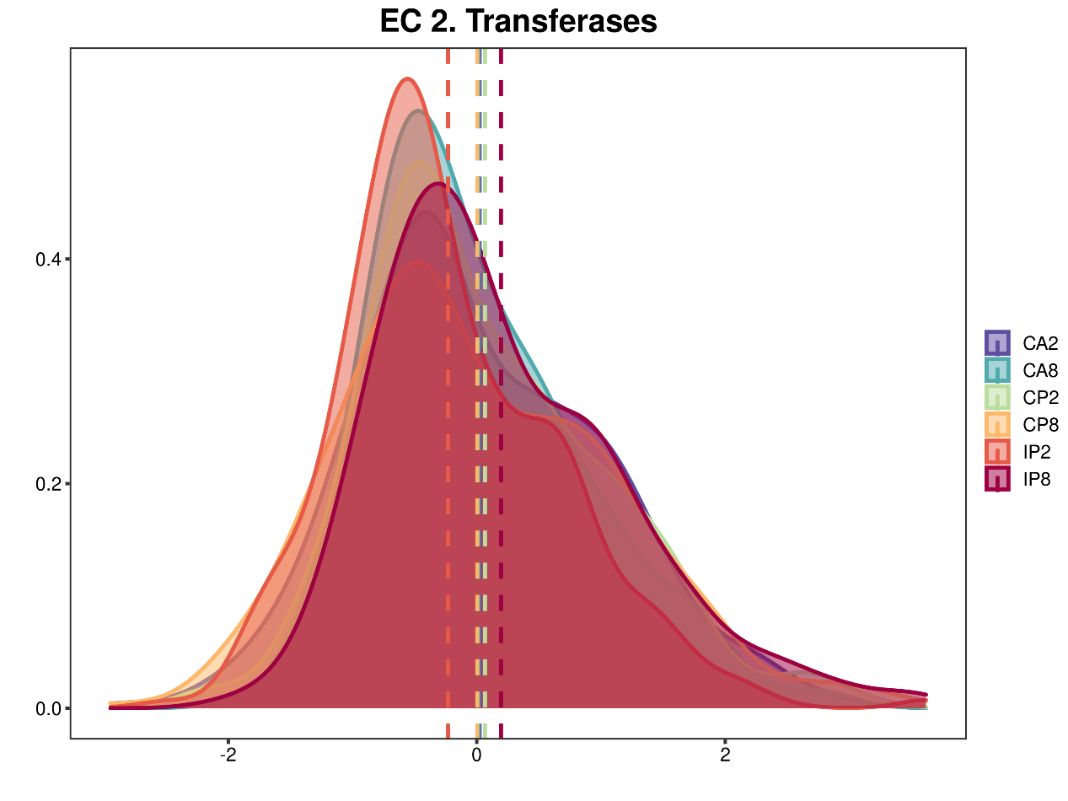

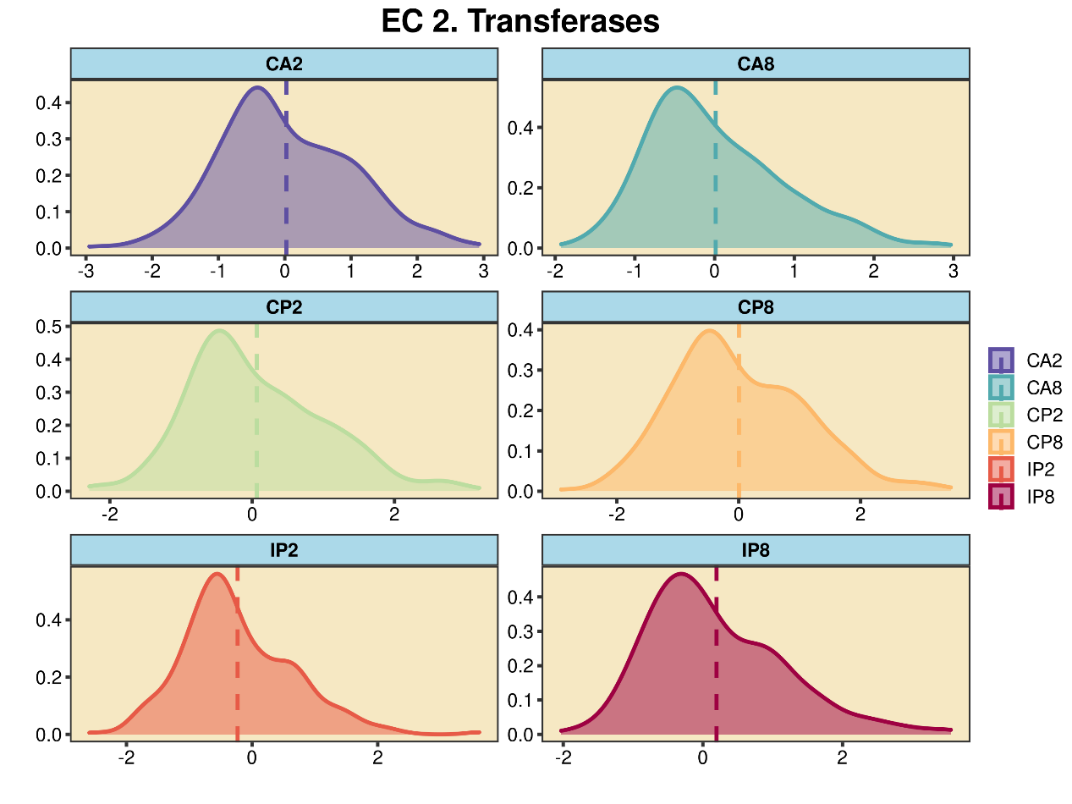

**Figure S2. Distribution and abundance of transferases among test conditions.**

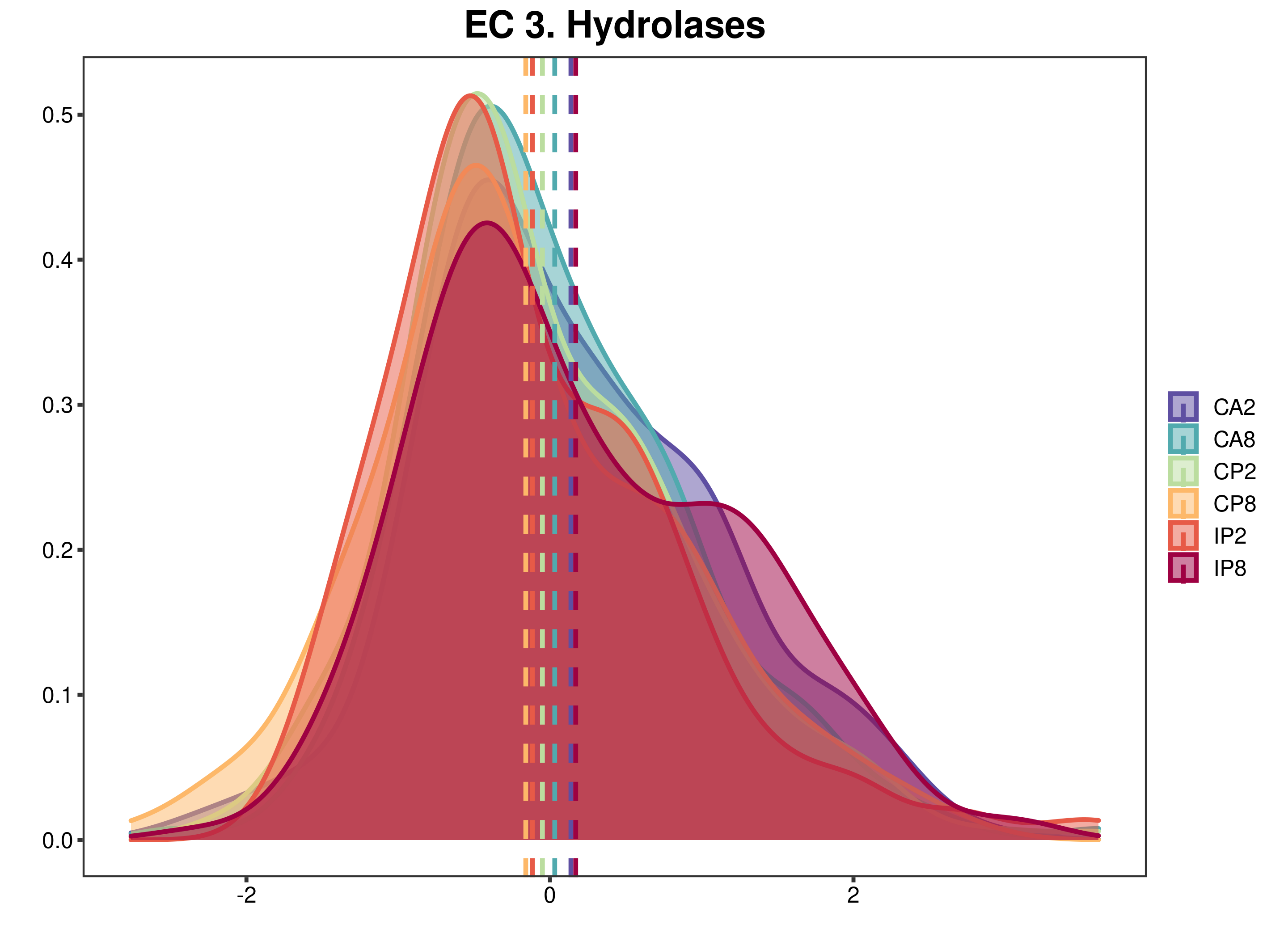

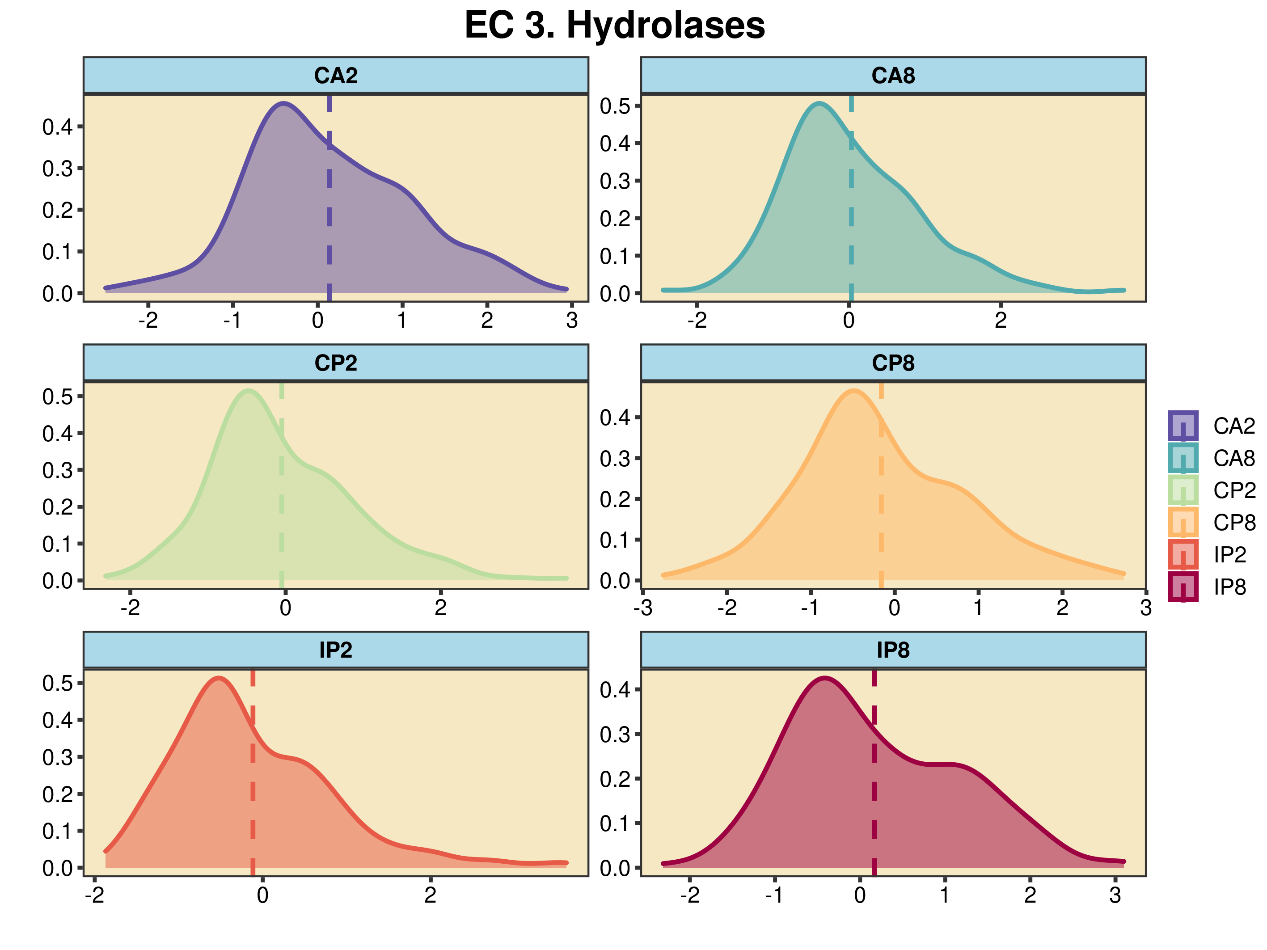

**Figure S3. Distribution and abundance of hydrolases among test conditions.**

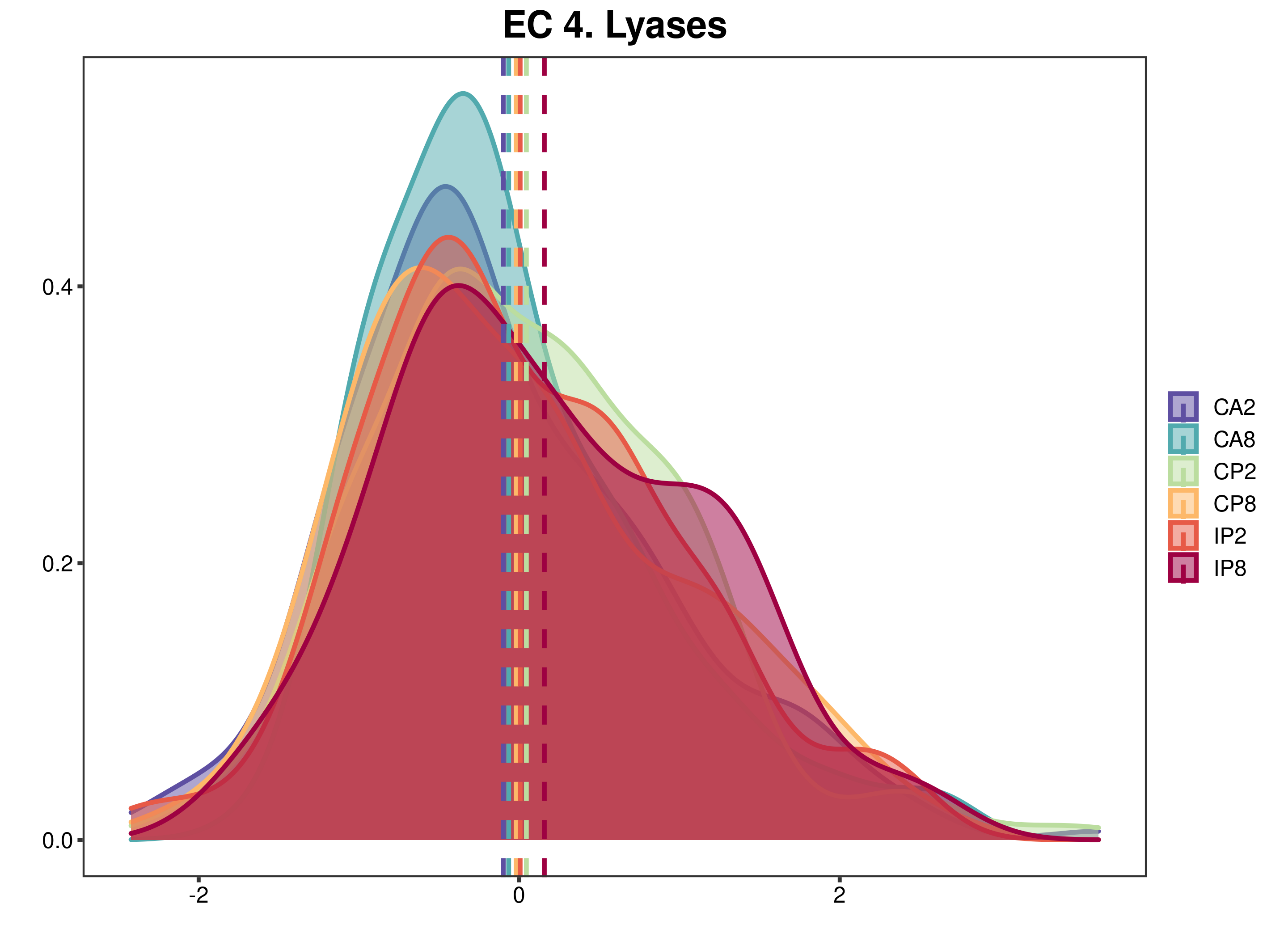

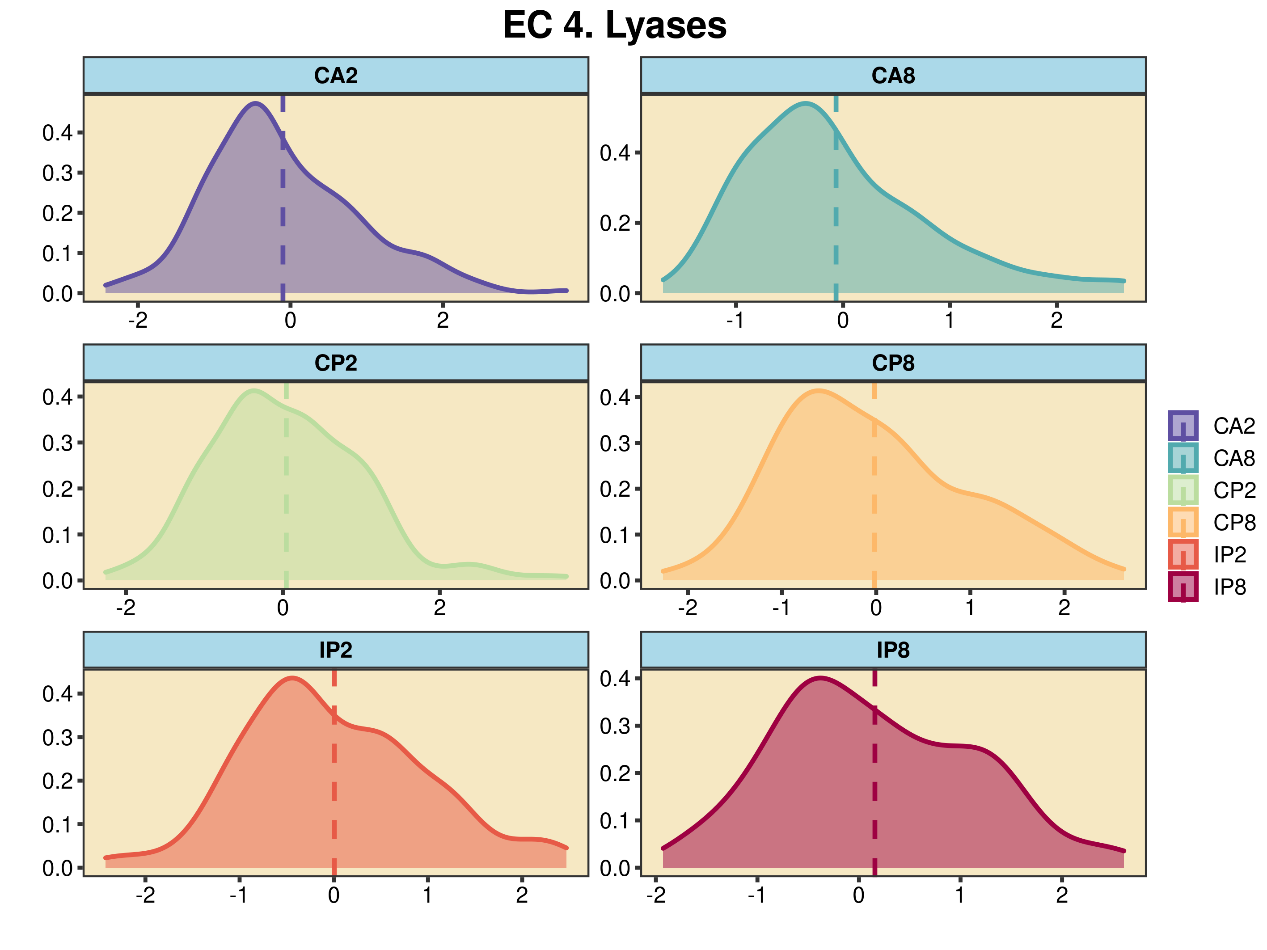

**Figure S4. Distribution and abundance of lyases among test conditions.**

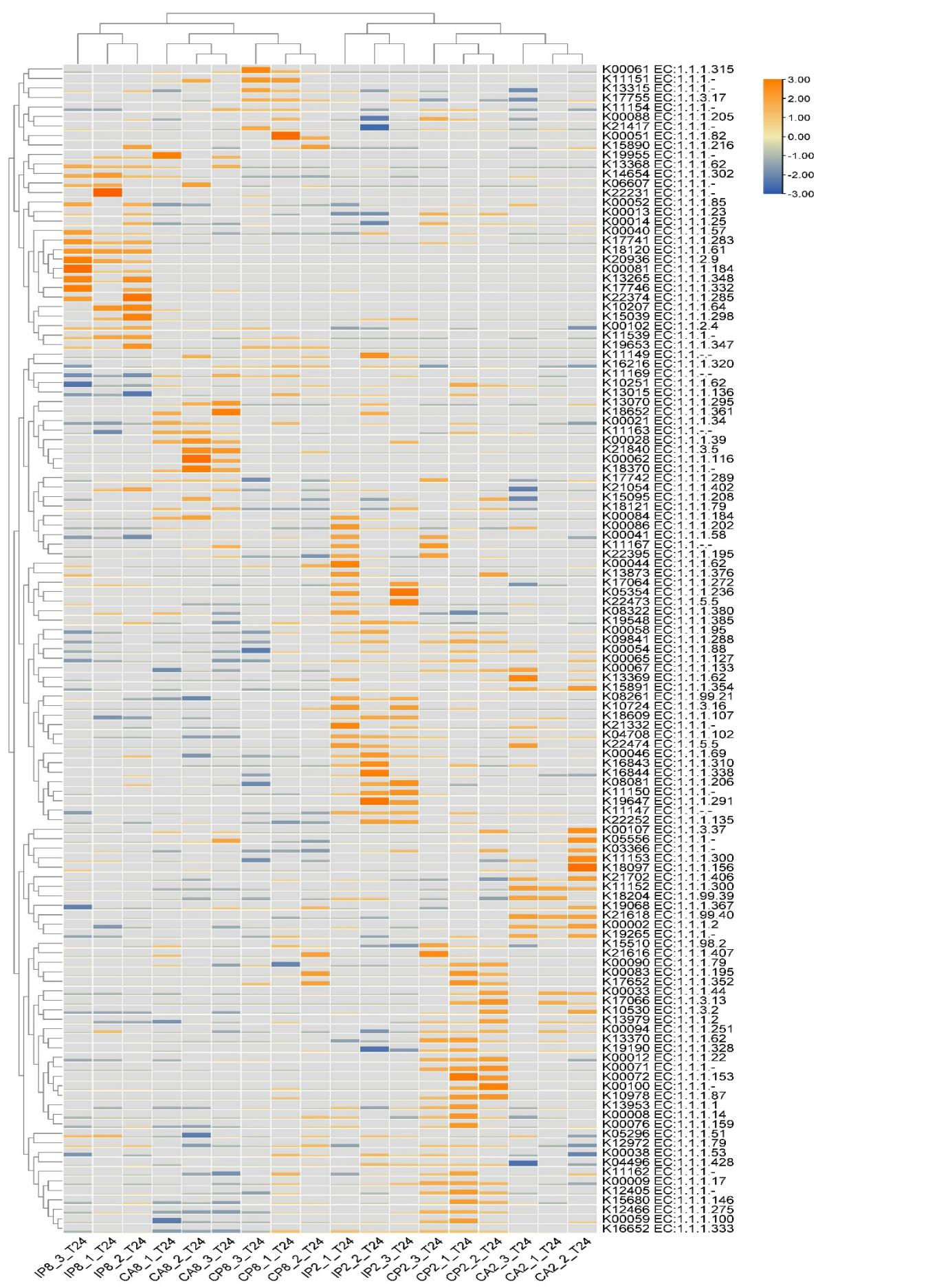

**Figure S5. Distribution and abundance of enzymes from EC1 among test conditions.**

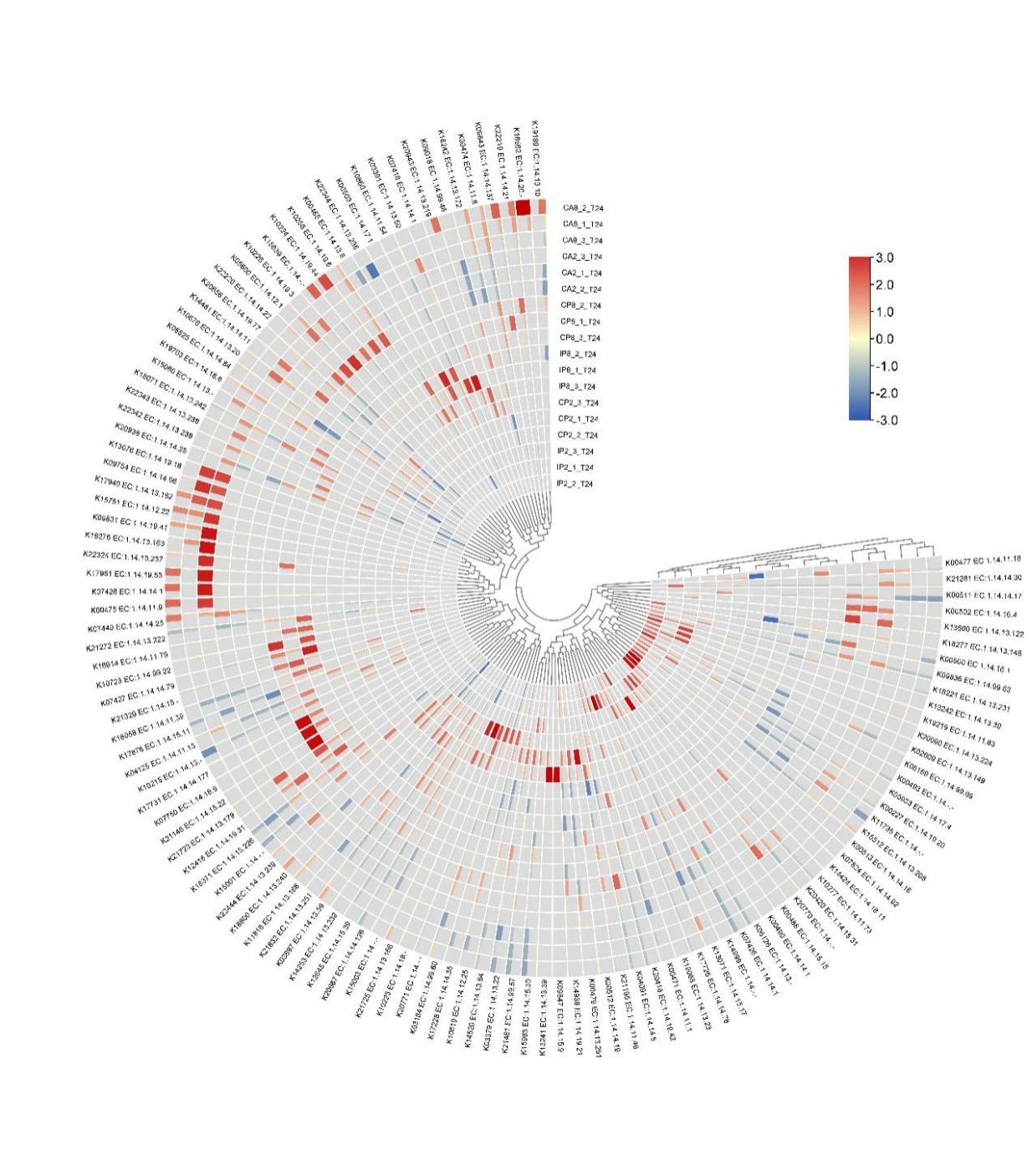

**Figure S6. Distribution and abundance of enzymes from EC1.14. among test conditions.**

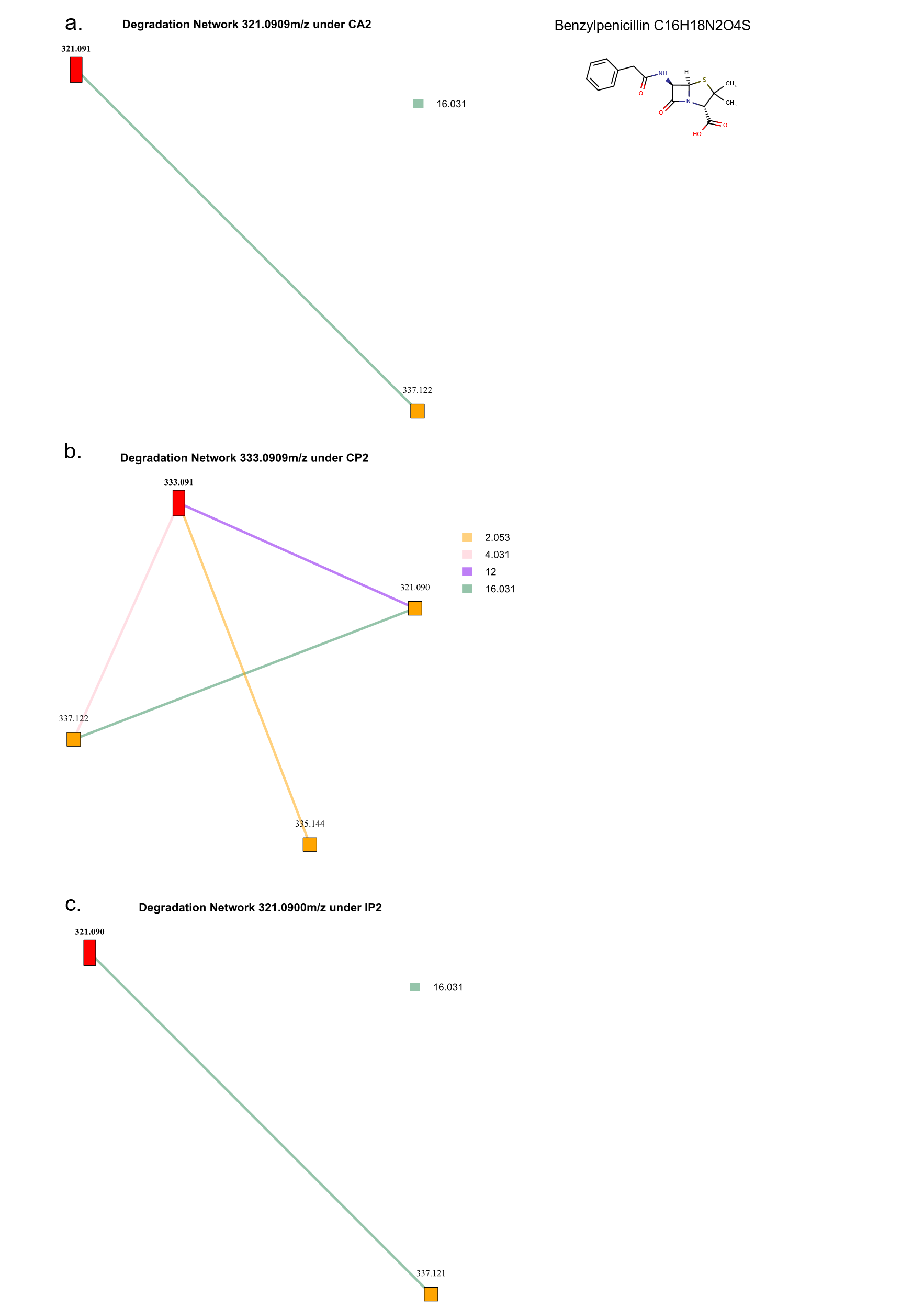

**Figure S7. Degradation pathway proposed by PMD analysis for** **Benzylpenicillin under conditions CA2 (a), CP2 (b), IP2 (c).**

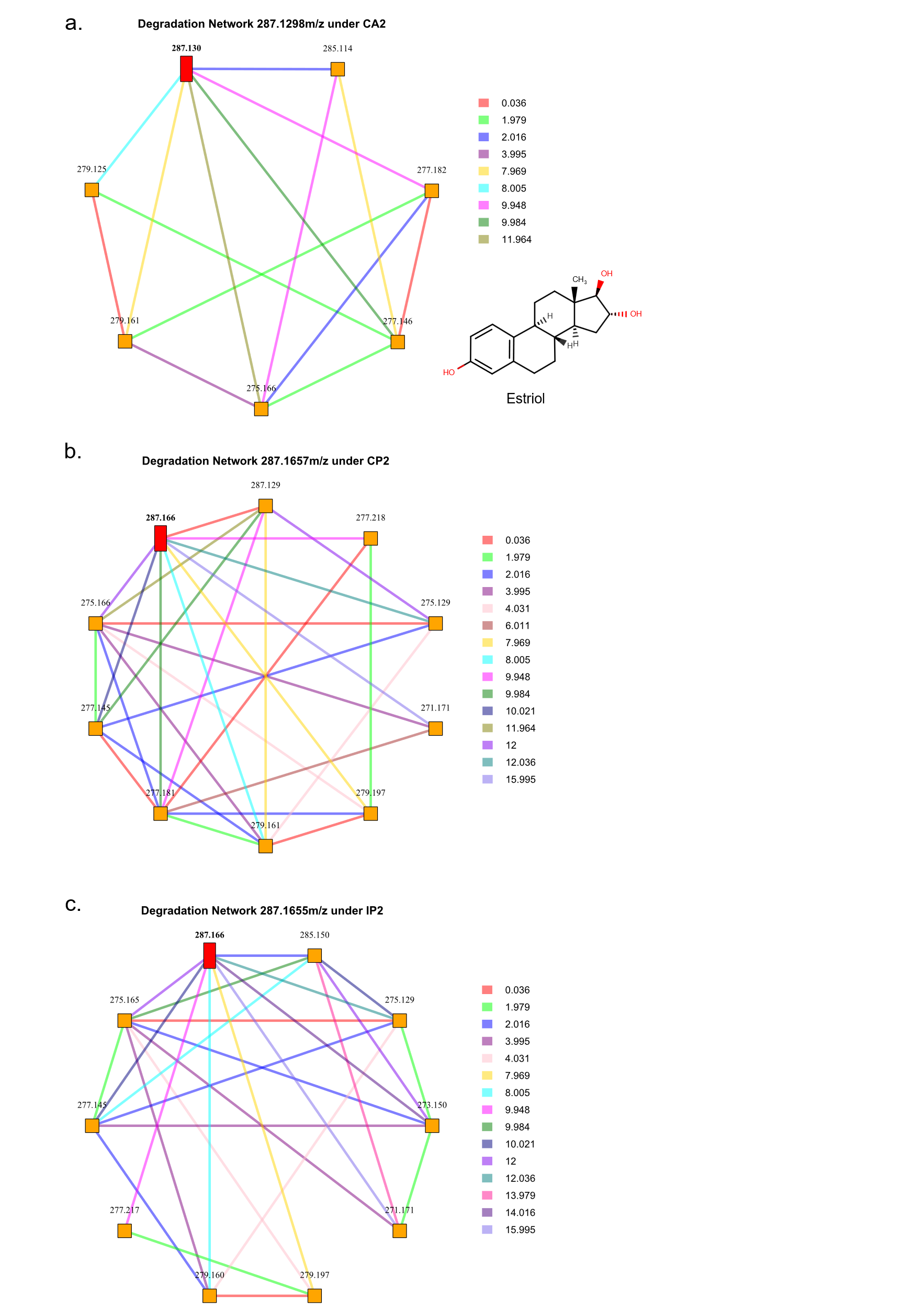

**Figure S8. Degradation pathway proposed by PMD analysis for Estriol under conditions CA2 (a), CP2 (b), IP2 (c).**

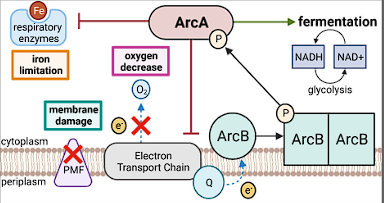

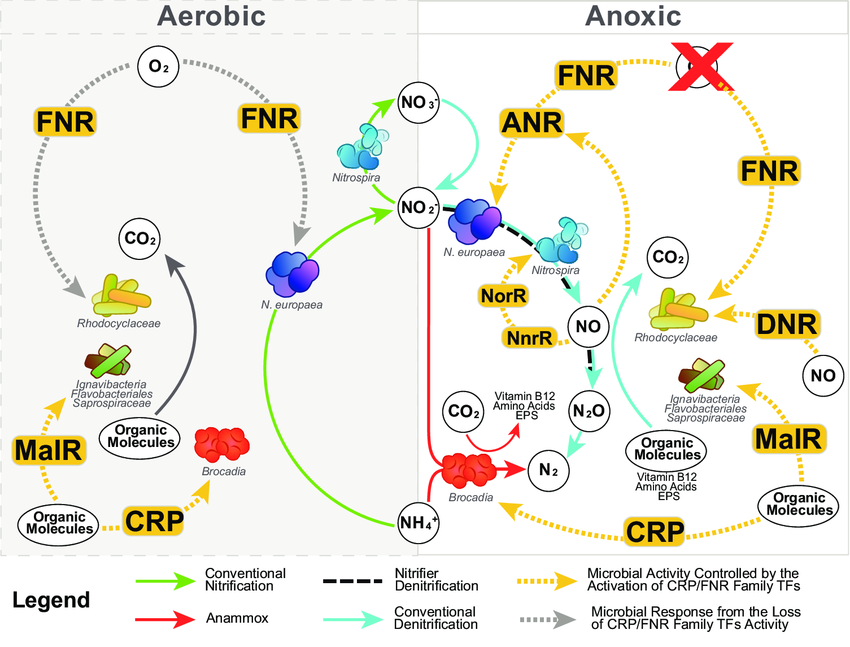

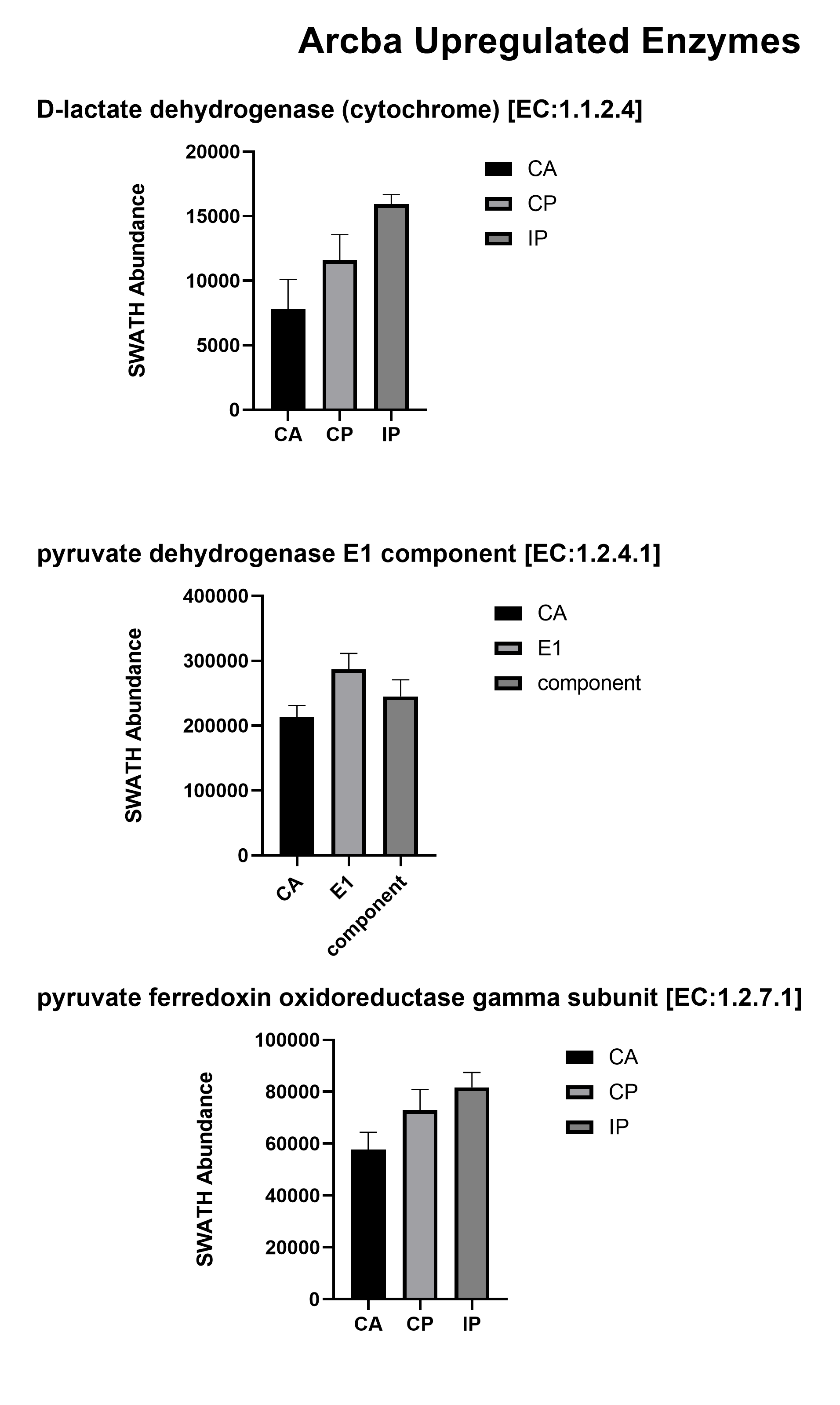

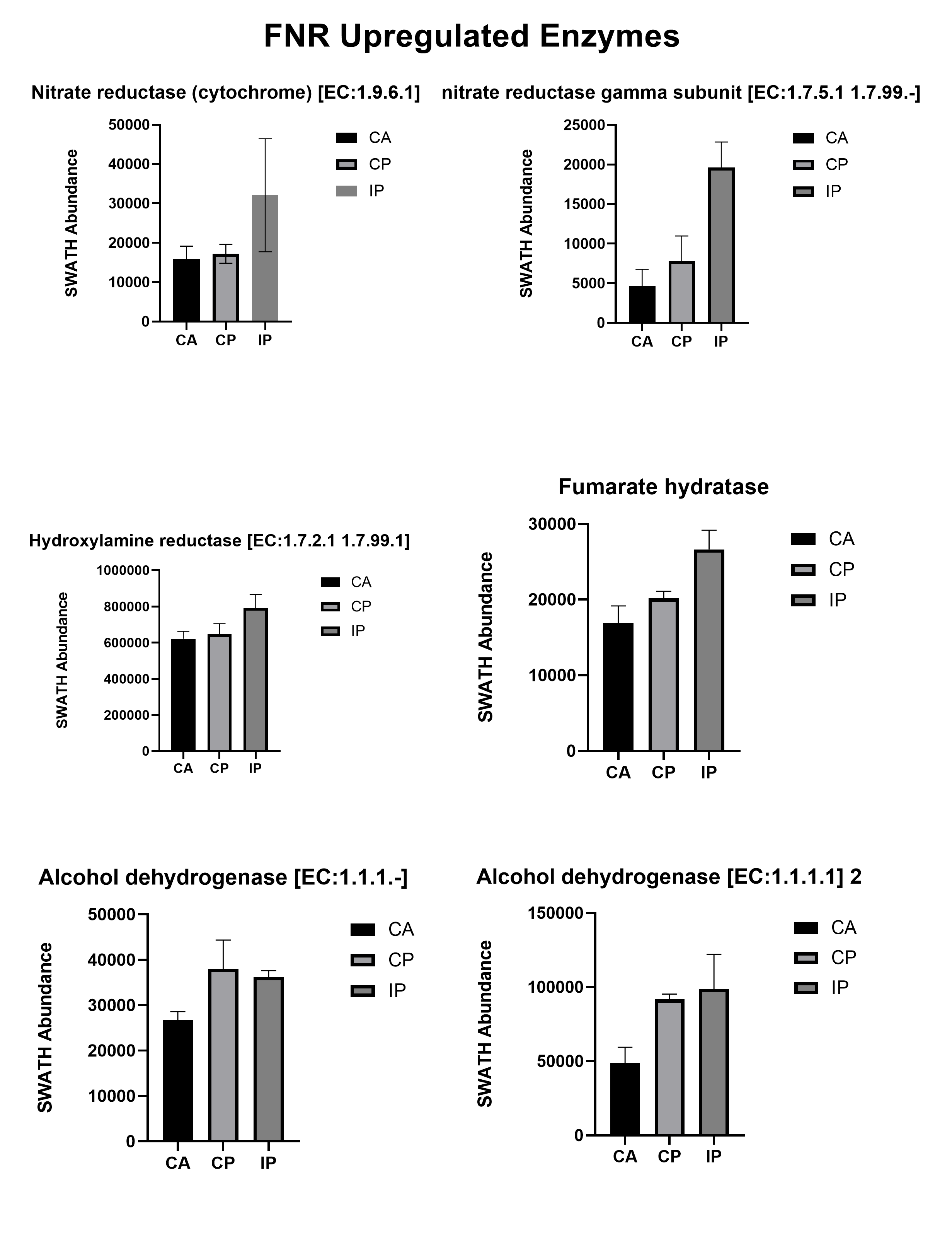
